# Leukocytes use endothelial membrane tunnels to extravasate the vasculature

**DOI:** 10.1101/2024.10.28.620560

**Authors:** Werner J. van der Meer, Abraham C.I. van Steen, Eike Mahlandt, Loïc Rolas, Haitao Wang, Janine J.G. Arts, Lanette Kempers, Max L.B. Grönloh, Rianne M. Schoon, Amber Driessen, Jos van Rijssel, Ingeborg Klaassen, Reinier O. Schlingemann, Yosif Manavski, Mark Hoogenboezem, Reinier A. Boon, Satya Khuon, Eric Wait, John Heddleston, Teng-Leong Chew, Martijn A. Nolte, Sussan Nourshargh, Joachim Goedhart, Jaap D. van Buul

**Affiliations:** Vascular Cell Biology Lab, Dept. Medical Biochemistry, Amsterdam UMC, University of Amsterdam; Molecular Cell Biology Lab, Dept. Molecular Hematology,Sanquin Research, and Landsteiner Laboratory, Amsterdam, the Netherlands; Leeuwenhoek Centre for Advanced Microscopy, section Molecular Cytology, Swammerdam Institute for Life Sciences, University of Amsterdam, Amsterdam, the Netherlands; Centre for Microvascular Research, William Harvey Research Institute, Faculty of Medicine and Dentistry, Queen Mary University of London, London, UK; Centre for Inflammation and Therapeutic Innovation, Queen Mary University of London, London, UK; Department of Ophthalmology at Amsterdam UMC, location University of Amsterdam; Department of Ophthalmology, University of Lausanne, Jules-Gonin Eye Hospital, Fondation Asile des Aveugles, Lausanne, Switzerland; The Institute for Cardiovascular Regeneration, Center of Molecular Medicine, Goethe University, Frankfurt, Germany; Core facility at Dept. Molecular Hematology, Sanquin Research and Landsteiner Laboratory, Amsterdam, the Netherlands; Advanced Imaging Center at Janelia Research Campus, Howard Hughes Medical Institute, Ashburn, VA, USA

**Keywords:** Transendothelial Migration, Transendothelial Migration Tunnel, Endothelium, Leukocyte, Neutrophil, Actin, Membrane, Inflammation, VE-cadherin, PECAM-1

## Abstract

Upon inflammation, leukocytes extravasate through endothelial cells. When they extravasate in a paracellular manner, it is generally accepted that neighbouring endothelial cells physically disconnect to open cell-cell junctions, allowing leukocytes to cross. When carefully examining endothelial junctions, we found a partial membrane overlap of endothelial cells beyond VE-cadherin distribution. These overlaps are regulated by actin polymerization and, although marked by, do not require PECAM-1, nor VE-cadherin. Neutrophils prefer wider membrane overlaps as exit sites. Detailed 3D analysis of endothelial membrane dynamics during paracellular neutrophil transmigration in real-time, at high spatiotemporal resolution using resonant confocal and lattice light-sheet imaging, revealed that overlapping endothelial membranes form a tunnel during neutrophil transmigration. These tunnels are formed by the neutrophil lifting the membrane of the upper endothelial cell while indenting and crawling over the membrane of the underlying endothelial cell. Our work shows that endothelial cells do not simply retract upon passage of neutrophils but provide membrane tunnels, allowing neutrophils to extravasate. This discovery defines the 3D multicellular architecture in which the paracellular transmigration of neutrophils occurs.

## Introduction

The immune system protects our body from bacteria, viruses, parasites, and injury. To reach the site of infection, immune cells extravasate the circulation and penetrate the vessel wall, in a process known as leukocyte transendothelial migration (TEM). This process is understood to occur in at least three distinguished steps: rolling, adhesion, and diapedesis. The latter step can occur in two ways: paracellular, i.e., through the cell-cell junctions, or transcellular, i.e., through the endothelial cell body^1^. The TEM model was first proposed by Butcher, and further defined by Springer^2,3^. Although many studies have added to our understanding, the principles of the “*multistep paradigm*” model still stand. One of the dogmas is that during the final diapedesis step, when the leukocytes penetrate the endothelium in a paracellular manner, neighboring endothelial cells detach. It is generally believed that the detachment of the two, or sometimes more, adjacent endothelial cells is triggered by the loss of trans-VE-cadherin interactions^4,5^, although VE-cadherin may also be redistributed, like a “trap-door” mechanism^6,7^.

Several previous structural studies focusing on the architecture of the vessel wall using transmission or scanning electron microscopy (EM) revealed that the endothelial cells do not simply lay side by side, like a sheet of epithelial cells, but in fact, partially overlap ^8–10^. It was observed in rat airways that the membrane from one endothelial cell protrudes underneath the membrane of the neighboring endothelial cell^11–13^. Despite this knowledge, this overlap between individual endothelial cells is often overlooked or ignored. Analyzing different vascular beds from a confetti-knock-in mouse model, we found these overlaps to be present in the vascular beds of the lung, liver, spleen, retina, and cremaster muscle. *In vitro* transmission EM images confirmed the overlaps in endothelial cell cultures. Using fluorescently membrane-labeled mosaic endothelial cell monolayers, we found that in a monolayer, virtually all endothelial cells overlap and function as preferred exit sites for leukocytes during inflammation. Real-time confocal and lattice light-sheet microscopy revealed that leukocytes squeezed themselves through the overlapping endothelial membranes using a so-called “transendothelial migration (TEM) tunnel”. Detailed analysis revealed that leukocytes lifted the membrane of the upper endothelial cell and crawl over and indented the membrane of the underlying endothelial cell, establishing a membrane tunnel through which the leukocytes squeezed. Differential dynamics of the junctional proteins VE-cadherin and PECAM-1 were found in this tunnel structure. Whereas VE-cadherin is excluded from the dissociated membranes and only present on the sides of the tunnel where the endothelial membranes are still connected, PECAM-1 was seen to decorate the entirety of the tunnel both *in vitro* and *in vivo*.

Together, we show here for the first time the true three-dimensional architecture of paracellular TEM events. Specifically, our findings reveal that endothelial cells do not simply dissociate from each other but rather form a membrane-based transmigration tunnel that allows leukocytes to migrate through. These findings enhance our understanding of the spatiotemporal profile of paracellular leukocyte transmigration and present novel means through which endothelial cellular structures support leukocytes in breaching the endothelial monolayer.

## Results

### Endothelial cell membranes overlap *in vivo* and *in vitro*

To examine the morphology of endothelial cells in a monolayer, we used transmission electron microscopy, and found that endothelial membranes of cultured endothelial cells (ECs) partially overlap at junction regions (Figure 1A), in so-called fork-like overlaps and single overlaps (Figure S1A). Despite the different morphological phenotypes, the overlaps showed an average width of around 4 µm. These data are in line with other studies that revealed the existence of EC overlaps *in vivo* in rat airways^11,12^. Using a fluorescent protein with a plasma membrane anchor (CaaX), we confirmed that one endothelial cell protruded beyond the VE-cadherin lining (Figure 1B). We found similar overlaps in human artery ECs (Figure S1B). To characterize the dynamics of the endothelial membrane overlap, two populations of endothelial cells were transduced with either mNeonGreen-CaaX or mScarlet-CaaX and a mosaic endothelial monolayer was generated (Figure 1C). Alternatively, we used mTurquoise2-CaaX and YFP-CaaX with similar results. For quantification, confocal Z-stacks were processed by thresholding the maximum projection of each color channel and quantifying the overlap between the two channels. The result is an overlap of 4-5 µm, comparable to what was measured with the electron microscope (Figure 1D). We additionally used a vessel-on-a-chip model^14^ to mimic human physiology more closely and found the presence of endothelial membrane overlaps (Figure S1C). To study whether endothelial membranes overlapped in different vascular beds *in vivo*, we used the Confetti^fl/wt^ Cdh5^CreERT2^ mice to induce endothelium-specific stochastic expression of membrane-bound CFP and cytosolic YFP/RFP^15^. We detected overlapping endothelial membranes in the lung, liver, and spleen (Figure 1E) as well as the cremaster muscle (<u>Video 1</u>). Moreover, we found overlapping endothelial membranes in inflamed retinas of Cynomolgus monkeys (Figure S1D)^16^. These data indicate that endothelial membrane overlaps can be found in different vascular beds.

**Figure 1:**
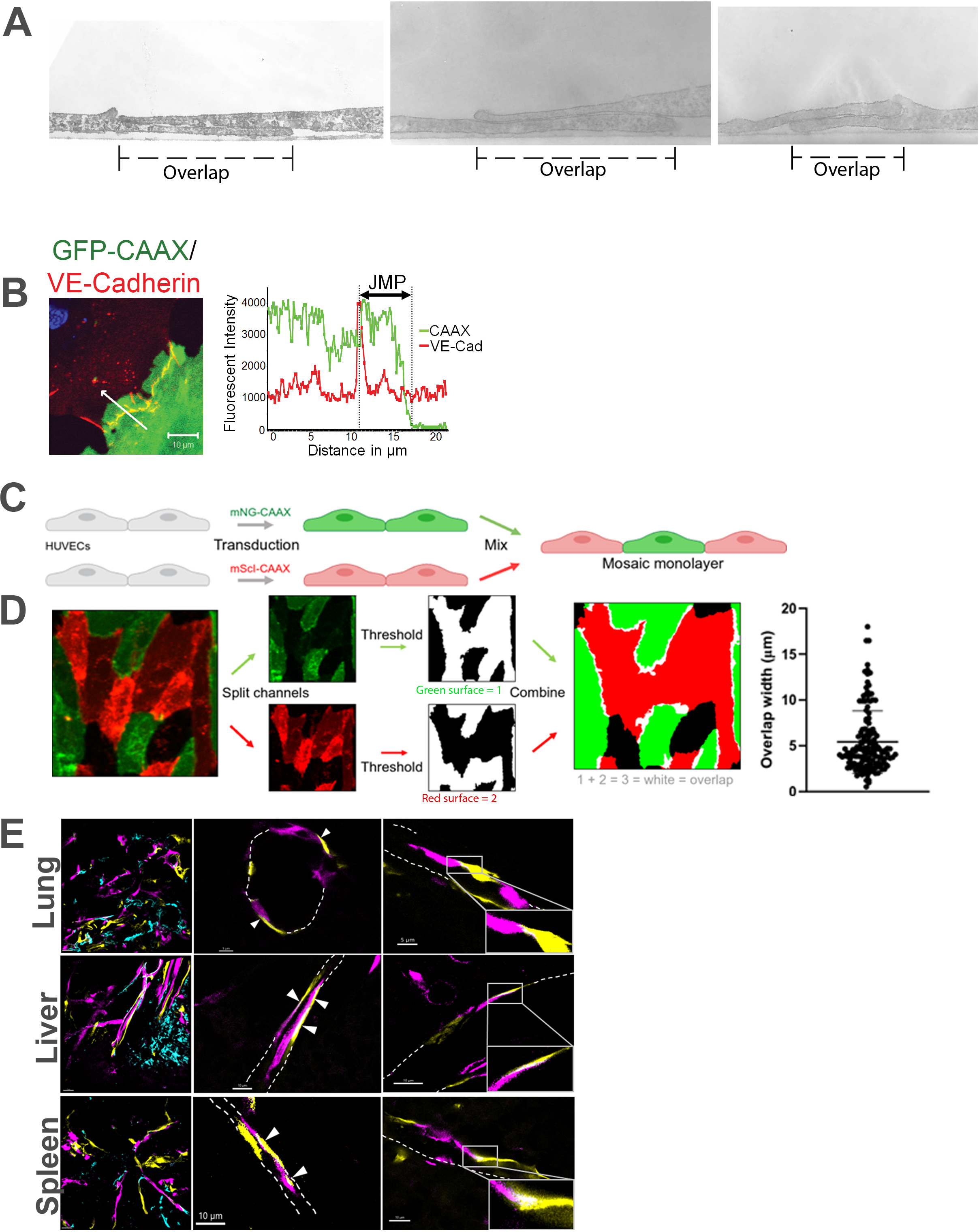
*Adjacent endothelial cell membranes overlap*: (**A**) Transmission electron microscopy is used to show overlapping endothelial membranes (dashed lines). Representative images shown of three separate experiments with 15 junctions imaged in total. Average junctional width of these overlaps is approximately 4 µm. (**B**) HUVEC transfected with membrane marker CAAX-GFP (green) and stained for VE-cadherin (red). Membrane continues beyond VE-cadherin stain. Representative image of three separate experiments with >10 junctions imaged per replicate. Graph on the right shows the intensity profile of the arrow with VE-cadherin in red and membrane in green. Overlap is marked as JMP (Junction Membrane Protrusion). Bar, 10 µm. (**C**) Schematic drawing of generation of mosaic endothelial monolayers. Mosaic monolayers were obtained by mixing two pools of HUVECs that were transduced with CAAX fluorescently labeled with either one of the color combinations mNeonGreen/mScarlet or YFP/mTurquoise2. (**D**) Maximum projections of confocal Z-stacks were split into separate channels. The resulting binary images were assigned different values, after which the images were summed to reveal overlapping surfaces. The mean overlap width of CAAX-expressing HUVECs is 5.4µm. n=142 junctions, 4 independent experiments. (**E**) Confetti^fl/wt^Cdh5^CreERT2^ mice were used to study EC overlap in the lung, liver, and spleen. Displayed are: Red fluorescent protein (Magenta), Yellow fluorescent protein (Yellow) and Cyan fluorescent protein (Cyan). Overlapping endothelial cells are detected in the different organs as indicated by arrows.

### Overlapping membranes depend on actin regulation and are favored for TEM

To investigate the dynamics of the endothelial overlap, we followed mosaic EC monolayers in time, quantified the overlap width, and found that the overlap remained stable at around 4 µm for at least 60 minutes (Figure 2A). To study whether the actin cytoskeleton is regulating EC overlap, we used small-molecule inhibitors. These inhibitors were administered to CaaX-expressing mosaic EC monolayers and the overlap dynamics were recorded in real-time. The results showed that blocking actin regulation reduced the overlap width, ultimately resulting in a loss of cell-cell contacts and gap formation (Figure 2B). Interestingly, blocking Rho-kinase also resulted in a drop in overlap region, indicating that both myosin-mediated actin contractility and regulation of actin turnover are necessary for the dynamics of endothelial membrane overlap.

**Figure 2:**
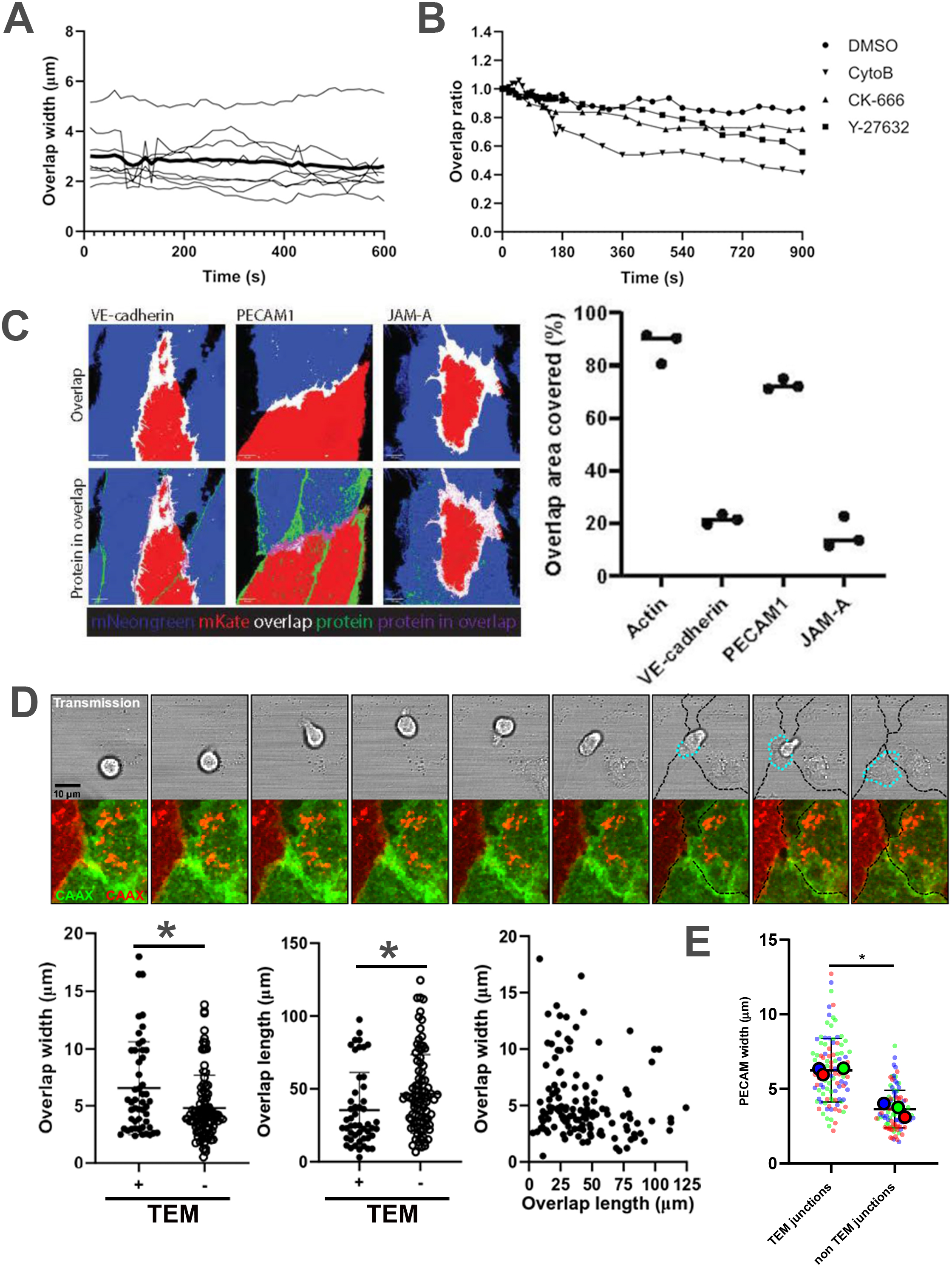
Characterisation of the endothelial overlap: (**A**) Endothelial overlap width of individual junctions and their average (thick line) quantified over time. N=9: 3junctions per time series in 3 separate experiments (**B**) Overlap width TNFa inflamed HUVECs quantified over time upon actin (CytoB or CK-666) or Rho-kinase (Y-27632) inhibition. Overlap quantified as a relative ratio to t0 pre-treatment, n>5 junctions per condition. **(C)** Quantification of junctional protein coverage of the overlap area shown in percentage. Data consists of 3 independent experiments, with 10-15 overlap areas per replicate. Dots represent the mean of each replicate. (**D**) Confocal microscopy images of a transmigrating neutrophil through mosaic HUVEC monolayer grown in an IBIDI flow slide. Upper panels: transmission images of one confocal Z-slice, lower panels: overlay of maximum projections of confocal Z-stacks of mScarlet- and mNeonGreen-Caax. Stills from time series, 40s apart. Overlap width of TEM event junctions (+,n=50) is significantly wider compared to non TEM event junctions (-,n=92) over 6 independent experiments (Mann Whitney, p=0.0242). The length of these TEM event junctions is significantly shorter compared to non TEM junctions (Mann Whitney, p=0.00037).Overlap length and width are negatively correlated (Pearson r=-0.18, p=0.0324), indicating that shorter overlapping areas are wider.. (**E**) HUVEC monolayers stimulated with TNFa for 4 hours, stained with non-blocking PECAM1 antibody, comparing PECAM width of TEM event junctions (n=102) versus non TEM event junctions (n=108). Colors indicate the three replicates, small dots represent individual junctions, big dots their average. PECAM width is significantly wider at TEM event junctions compared to non TEM event junctions (Unpaired t-test, p=0.001).

Next, we investigated the distribution of junctional proteins covering the EC overlap. We found that PECAM-1 covered overlapping areas for more than 70% whereas VE-cadherin and JAM-A covered the overlap for only 15% (Figure 2C). However, when PECAM-1 was depleted, we observed no change in overlap width (Figure S2A), indicating that PECAM-1 marked the overlaps but was not required for their formation. Treating EC monolayers with TNFα induced an elongated EC morphology, an established inflammatory phenotype^17^ but did not influence the dynamics or width of the overlaps (Figure S2B). Remarkably, when focusing on TEM events, we found that neutrophils preferred to cross EC junctions at areas with wider overlaps (Figure 2D; left panel). Using a non-blocking PECAM-1 antibody it was found that junctions with wider PECAM-1 distribution were preferred as well for neutrophil TEM (Figure 2E), in line with the fact that PECAM-1 covered over 70% of the membrane overlap area (Figure 2C). Moreover, junctions that supported TEM were significantly shorter in length than non-TEM junctions within the same monolayer (Figure 2D and S2C). Interestingly, endothelial membrane overlap width and length were slightly, yet significantly, inversely correlated (Figure 2D). These results indicate that the shorter sides of elongated inflamed endothelial cells overlapped more. It has been suggested that tricellular junctions may act as TEM hotspots for neutrophils when crossing endothelial monolayers^18,19^. We found that neutrophils crawled on average 15 µm before crossing a junction (Figure S2D). Almost all neutrophils crossed the first junction they encountered, and some preference for tricellular junctions was detected, although the difference was not significant (Figure S2D).

### Overlapping membranes form a tunnel to allow paracellular diapedesis

To accurately capture all structural changes of both cell types, i.e., the leukocyte and the endothelial cell during the extravasation process, it is required to use extremely fast data acquisition at a high-resolution level. We used confocal resonant scanning to increase the acquisition speed in 3D significantly^20^ and applied volume rendering software analysis on the raw data to increase visibility and interpretation of the data (Figure S3A). We found that the overlapping endothelial membranes formed a tunnel-like structure as the neutrophils crossed (Figure 3A left panel, Video 2 and 3). All TEM events that were detected, showed the presence of such TEM tunnels (Figure 3A, right panel). The average diameter of nine tunnels imaged over three separate experiments was of 5.5 µm, in line with the gap size that has been measured in 2D TEM assays by us and others *in vitro* ^5,11,21,22^, as well as *in vivo* ^23^. Moreover, we observed that endothelial cells were not dissociating from each other but rather were lifted by the neutrophil to form the TEM tunnel (Figure 3B and Video 4).

**Figure 3:**
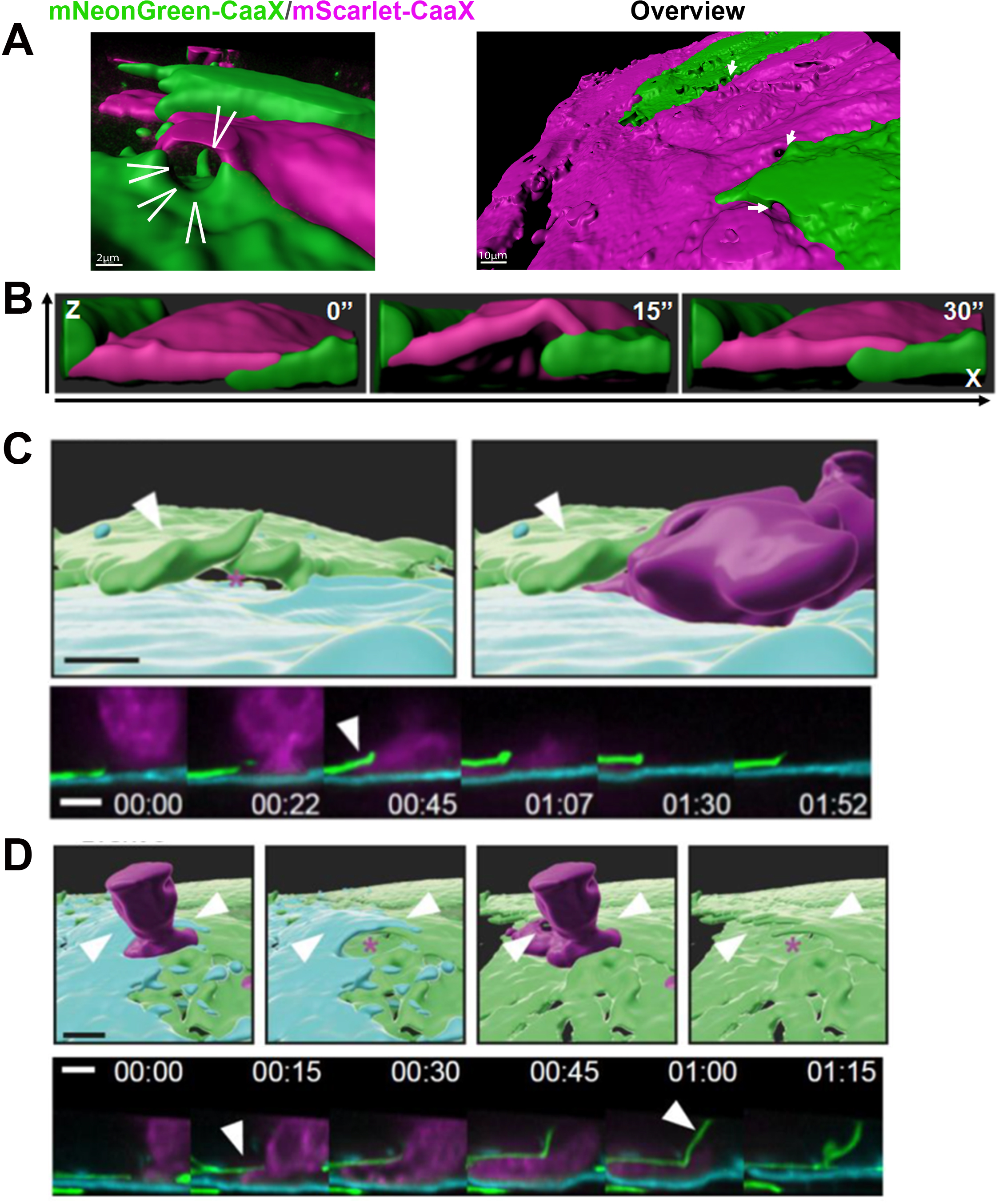
Overlapping endothelial membranes form a tunnel structure upon extravasation of neutrophils: (**A**) Surface rendered still from confocal microscopy movie showing multiple tunnel structures (white arrows) formed at the moment of neutrophil (not shown) diapedesis through a monolayer transduced with mScarlet-CaaX and mNeonGreen-CaaX, scale bar=10μm. Zoom shows transduced cell (magenta) forming the top and the mNeonGreen-CaaX transduced cell forming the bottom of a tunnel, scale bar = 2 μm. Representative stills of 9 events from three independent experiments. **(B)** Orthogonal slice-view of tunnel formation during diapedesis of neutrophil (not shown).(**C**) Representative still (26 events captured) from a lattice light sheet microscopy movie showing tunnel formed by underlying (turquoise) and upper (green) endothelial cell as a neutrophil (magenta) extravasates), with orthogonal slice showing the non-rendered signal (**D)** Still from a lattice light sheet microscopy movie showing tunnel formed by underlying (green) and upper (turquoise) endothelial cell as neutrophil extravasates. Orthogonal slice view shows the non-rendered signal

To investigate these endothelial membrane structures during TEM with the highest possible spatial and temporal resolution, data was acquired using lattice light sheet microscopy (LLSM)^24^. In comparison to the point illumination of the confocal microscope, LLSM illuminates an entire sheet. The sheet is applied at an angle relative to the sample and can be moved to cover the entire sample (Figure S3B). This type of illumination allows fast scanning through the entire sample providing detailed spatial information in x, y, and z direction, with low phototoxicity. Endothelial cells stably expressed one of the CaaX-membrane markers and were grown into a mosaic-like monolayer. Data processing involves an automated deskew and deconvolution step^24^ (Figure S3C). The volume-rendered data showed the cell surface in great spatial detail, especially the overlapping endothelial cell membranes (Figure S3D). These membrane structures were difficult to capture using regular confocal microscopy as the LLSM obtains a Z-resolution of around 300 nm, depending on the mode used^24^, while confocal microscopy typically has a limit of 500 nm in Z-direction^25^.

The recordings showed that as a neutrophil arrived on an endothelial cell that displayed a membrane protrusion underneath its neighboring cell, the overlapping top EC membrane protrusion was lifted, facilitating the neutrophil migration underneath the upper EC (Figure 3C). This occurred rapidly within 2 minutes during which both endothelial cells covered the migrating neutrophil (Figure 3D).

For all events observed, at the basolateral level, an endothelial sheet was presented to the transmigrating neutrophil (Figure 4A; <u>Video 5</u>). The pore exclusively appeared when the neutrophil was present between the ECs, indicating that the neutrophil initiated the pore opening. The endothelium immediately started closing the pore once the neutrophil had passed (Figure 4B; <u>Video 5</u>). Interestingly, it was observed that both the top and the bottom EC contributed to the formation of the TEM tunnel. The bottom EC showed protrusions that acted as pillars and were attached to the top EC that formed the ceiling of the tunnel (Figure 4C, Top panel: Video5, bottom panel: <u>Video 6</u>). When further analyzing the structure inside the tunnel, we found that the bottom EC contributed to the walls of the tunnels by the formation of these pillar-like structures (Figure 4D, top panel: <u>Video 7</u>, bottom panel: <u>Video 8</u>). Although it is difficult to analyze in even greater detail, these structures may include focal adhesion sites, which we have shown are physical obstacles to neutrophils^26^.

**Figure 4:**
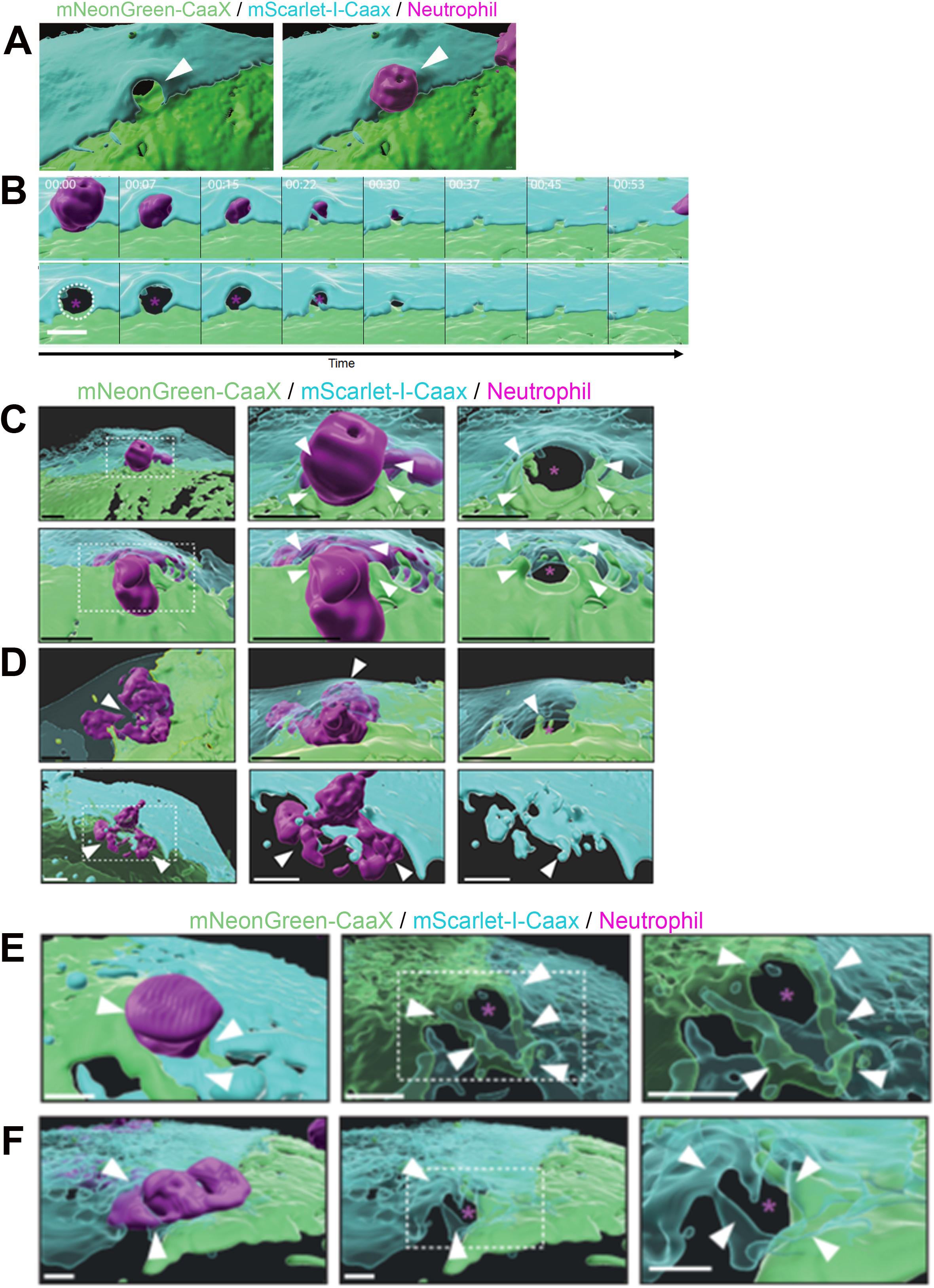
TEM tunnel dynamics and architecture (**A**) Surface rendering of endothelial membranes (turquoise, green) forming a TEM tunnel at the moment a neutrophil (magenta) is in mid-diapedesis (**B**) Surface rendering of endothelial membranes (turquoise, green) and their temporal dynamics during the closing of TEM tunnel **(C)** Surface rendering of endothelial membranes (turquoise, green) forming a TEM tunnel as a neutrophil (magenta) moves through imaged using lattice light sheet microscopy. **(D)** Endothelial membranes (turquoise, green) during opening and closing of TEM tunnel as neutrophil (magenta) moves through. Top panel shows the neutrophil extravasating, bottom panel visualizes only the endothelial cells. (**E,F**) Paracellular-transcellular TEM events show aberrant membrane configuration. Two examples of transmigration events that cannot be classified as either para- or transcellular diapedesis with endothelial membranes (turquoise, green) and transmigrating neutrophil (magenta). These events show tunnels with a more complicated architecture compared to the “simple” bottom and top cell configuration of TEM tunnels classified as paracellular. 26 TEM events were captured using the LLSM technique.

Previously, our lab showed that endothelial junctional membrane protrusions (JMPs) are linked to neutrophil TEM hotspots^27,28^. Therefore, we concentrated in more detail on these structures before, during, and after diapedesis. Some protrusions at diapedesis sites were characterized as JMPs (Figure S4A, <u>Video 9</u>), while other protrusions at diapedesis sites showed a more finger-like or filopodia appearance (Figure S4B, <u>Video 10</u>). Additionally, we observed the combination of both structures (Figure S4C, D: <u>Video 11</u>, E: <u>Video 12</u>). In every case, diapedesis events were accompanied by active membrane structures that originated from the endothelial cells.

### Diapedesis events can combine the para- and transcellular route

Already in the nineties, Butcher and Springer described several well-defined stages of transmigration including rolling, adhesion, crawling, and diapedesis^2,3^. They discriminated between the transcellular and paracellular diapedesis routes. Interestingly, we observed several events that cannot be categorized in either of the two routes. In some cases, diapedesis appeared close to a junction but not at a junction. Yet, both endothelial cells contributed by inducing dynamic membrane structures. whereas the tunnel pore seems to be made by only one of the two endothelial cells (Figure 4E, <u>Video 13</u>, F).

### Junctional protein dynamics in the TEM tunnel

As most neutrophils prefer the paracellular route to cross the endothelium, and as neutrophils made their way through an endothelial tunnel when transmigrating, we questioned how the proteins involved in leukocyte TEM were distributed during this event. We found that ICAM-1 was distributed along the tunnel, supporting the transmigration route of the neutrophil (Figure 5A), following previous work^29^. These data were recorded with a regular confocal microscope. Moreover, we found that PECAM-1 was distributed in a ring-like structure around the neutrophil when starting to penetrate (Figure 5B and S5A). This is in line with other studies ^30–32^. Using Airyscan imaging, we found PECAM-1 to decorate the inside of the TEM tunnel, suggestive of support during the diapedesis step (Figure 5C and <u>Video 14</u>). For VE-cadherin, we noticed that it was locally dispersed and only present at the sides of the tunnel (Figure S5B), in line with previous work from us and others^7,33,34^. By using 2 different FP-tagged VE-cadherin constructs, each transfected in adjacent endothelial cells, we generated a mosaic monolayer of fluorescently tagged VE-cadherin-positive endothelial cell-cell junctions and confirmed local dispersion of VE-cadherin at sites of diapedesis. Interestingly, when leukocytes penetrated the junction we found VE-cadherin to remain co-localized adjacent to diapedesis sites. We did not detect single-colored VE-cadherin, indicating that the trans-VE-cadherin interactions were maintained during diapedesis, at least at the edge of the TEM tunnels (Figure 5D and S5C). No single-colored VE-cadherin was detected at the TEM tunnel edges, indicating that VE-cadherin as a trans-interacting complex is pushed aside rather than a loss of VE-cadherin trans-interactions (Figure 5E). To show that single-colored VE-cadherin can be detected in this setup, endothelial cells were treated with EGTA to chelate calcium, as VE-cadherin trans-interactions depend on calcium. As a result, VE-cadherin-trans interactions dissociate into single-colored ones, whereas short-term thrombin treatment did not (Figure S5D, <u>Video 15</u>). These data suggest that the lateral membrane mobility of VE-cadherin, rather than its ability to dissociate, is a prerequisite for efficient leukocyte paracellular TEM.

**Figure 5:**
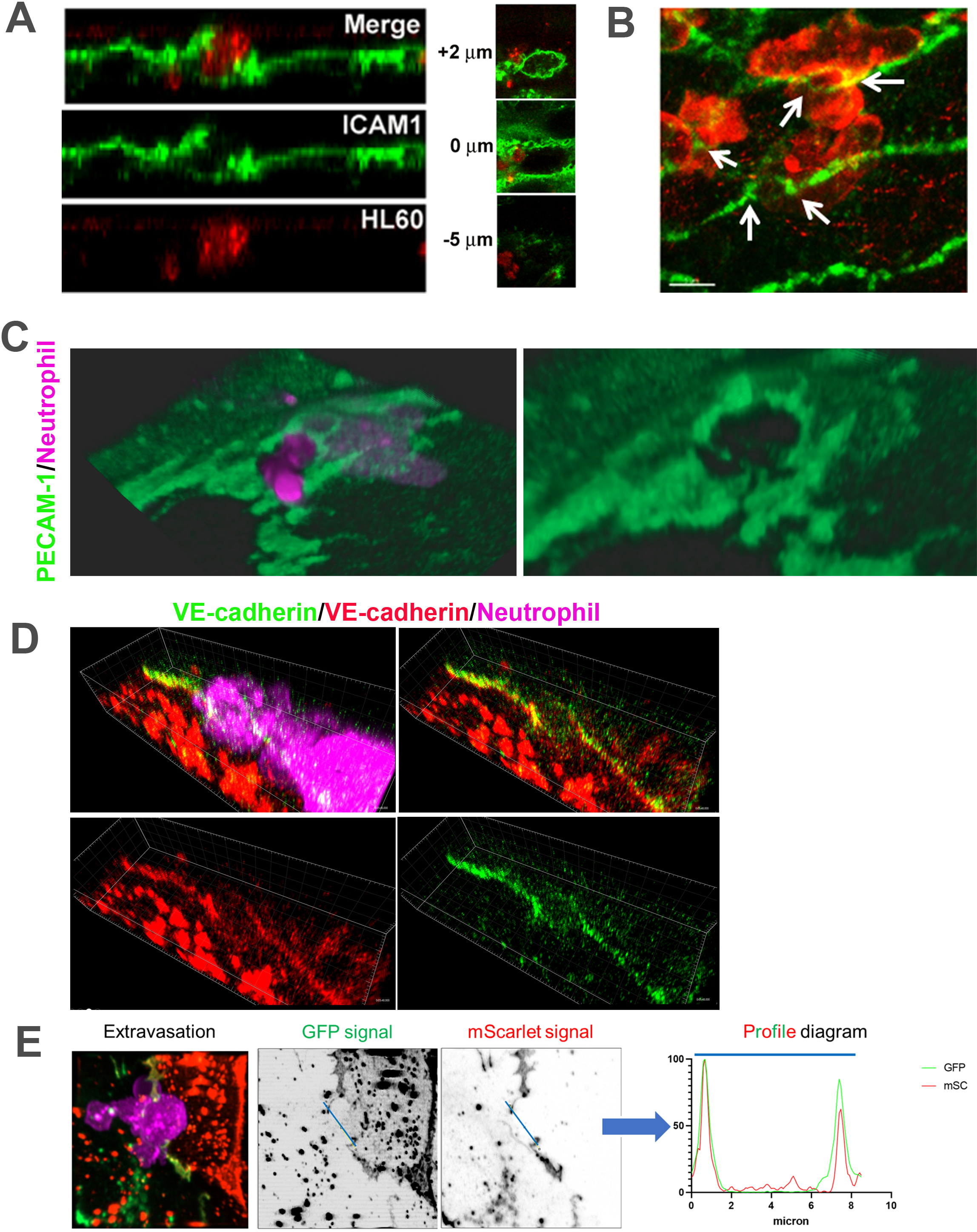
Distribution of junctional proteins in TEM tunnel (**A,B**) ICAM1 is shown to decorate both the top as the bottom cell of a TEM tunnel induced by a transmigrating HL60 cell. Representative image of 10 events from two separate experiments. (**C**) Representative still from a confocal movie of a neutrophil (magenta) transmigrating through a PECAM1-mNeonGreen (green) expressing HUVEC monolayer. Two independent experiments, (**D**) Still from a movie of a neutrophil (magenta) transmigrating through an endothelial junction with the upper right cell expressing VEcadherin-GFP (green), the lower left VEcadherin-mScarlet (red). (**E**) Representative (three independent experiments, 9 events) fixed transmigration event showing a neutrophil (magenta) during the diapedesis step with the right endothelial cell expressing VEcadherin-mScarlet (red), the left endothelial cell expressing VEcadherin-GFP (green). The fluorescent signal of both FP’s was plotted on a line drawn across this transmigration event, showing the signal of both FP’s on the sides of the TEM tunnel directly on top of each other.

### TEM tunnels decorated by PECAM-1 are observed *in vivo*

To study the TEM tunnels upon inflammation *in vivo*, we analysed neutrophil diapedesis in inflamed murine cremaster muscles, as observed using a 4D imaging platform with advanced spatial and temporal resolution^32^. Cremaster muscles were inflamed via transient induction of ischemia of the testes followed by restoration of blood flow and recording of neutrophil/vessel interactions in postcapillary venules according to Barkaway and Rolas et al^37^. Breaching of venular walls was investigated in real-time using neutrophil reporter mice *Lyz2-EGFP-ki* (display GFP^bright^ neutrophils) and following staining of endothelial cell-cell junctions by locally administered non-blocking anti-PECAM-1 antibodies^32^. Neutrophil extravasation was captured in real-time (Figure 6A and <u>Video 16</u>). To analyze TEM events in more detail, we used Imaris software to generate an iso-surface of PECAM-1 staining. We found neutrophils to cross the endothelial layer through thick PECAM-1-positive regions, similar to the observations detected using *in vitro* models (Figure 6B). The presence of PECAM-1 around the entire extravasating neutrophil indicated the presence of endothelial membranes both above and below the migrating neutrophil (Figure 6C and Video 17). Collectively, these results support the concept that neutrophils cross the endothelial monolayer by using membrane-based tunnels through two layers of adjacent endothelial cells.

**Figure 6:**
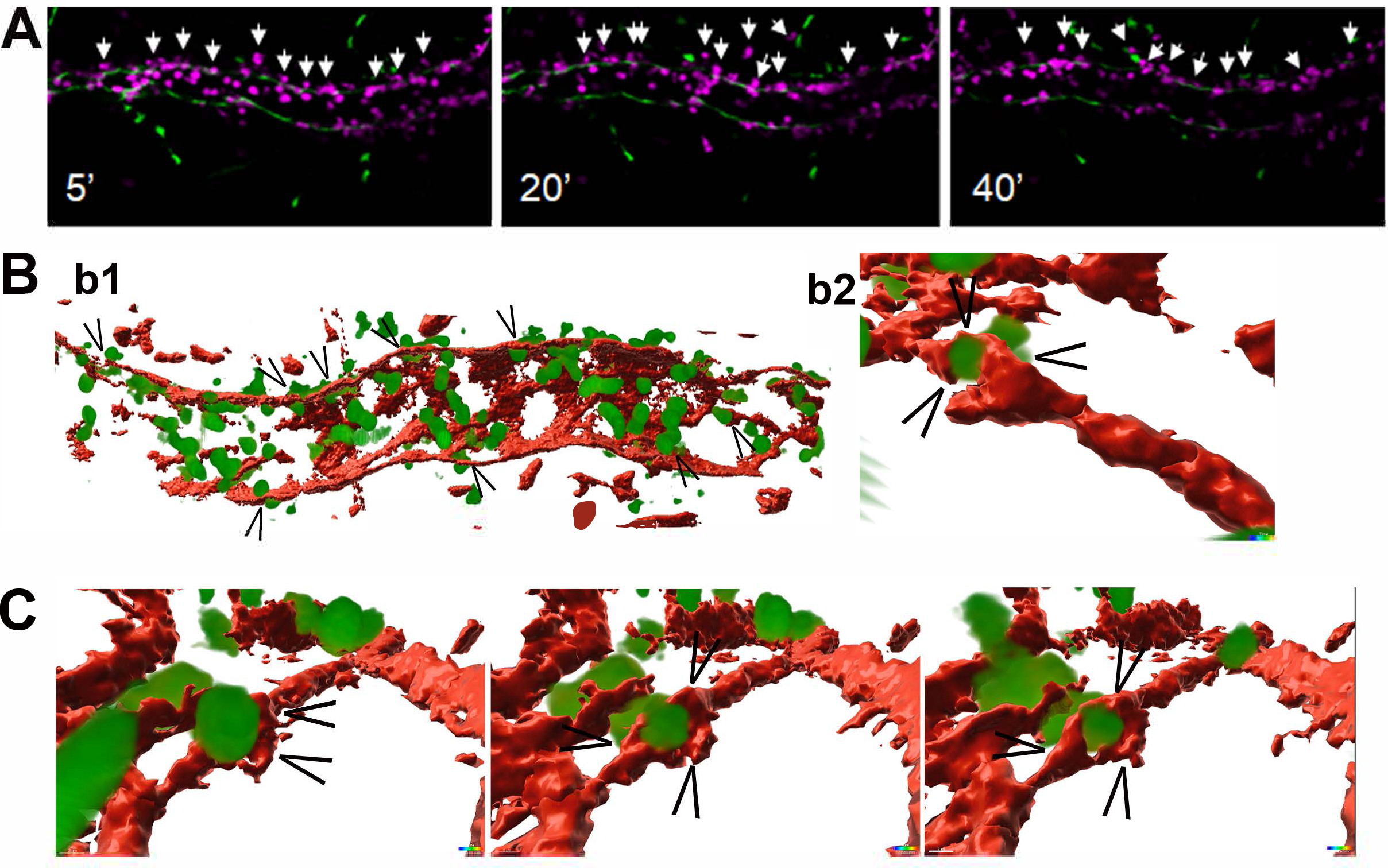
Spatiotemporal dynamics of PECAM-1 during leukocyte extravasation in vivo: Cremasteric venules of *LysM-EGFP-ki* mice (exhibiting GFP^high^ neutrophils) were immunostained *in vivo* for EC junctions with intrascrotal (i.s.) injection of Alexa Fluor-555-labeled anti-PECAM-1 mAb (PECAM-555; 4μg/mouse; shown in red). Inflammation was induced by transient ischemia of the testes (40 minutes), followed by blood flow restoration and recording of neutrophil/vessel interactions in postcapillary venules. (**A**) Panels show images of a postcapillary venular segment post reperfusion at different times (t=5’, 20’, and 40’ post-reperfusion), showing the development of an inflammatory response with neutrophils in magenta and PECAM-1 in green. White arrows indicate extravasating leukocytes. (**B/C**) Neutrophil (green) extravasation through endothelial junctions labeled for PECAM-1 (red) show presence of PECAM all around the transmigrating leukocyte. Images are representative of at least 6 independent experiments involving 6 mice.

## Discussion

During inflammation, circulating leukocytes cross the inflamed endothelial monolayer in a process called transendothelial migration (TEM). Neutrophils and monocytes mainly cross the endothelium in a paracellular manner, i.e., using the junctional route. It is the dogma in the field that adjacent endothelial cells transiently physically dissociate to allow leukocytes to pass and that endothelial cell borders start and stop at VE-cadherin localization^4,7^. However, we and others never observed that endothelial membranes retracted upon leukocyte extravasation; instead, tight gaps, supported by rings of tensile F-actin bundles, are induced between two adjacent endothelial cells to quickly close after leukocyte passage^21,22,32^. Moreover, here we show that adjacent endothelial cell membranes overlap, extending several micrometers beyond VE-cadherin, and that these overlapping membrane areas are favored by neutrophils to cross the endothelial barrier. The finding that endothelial cell membranes overlap is not new, in fact it has been recognized already for years^8–10^. The McDonald lab documented this phenomenon *in vivo* using scanning electron microscopy showing that endothelial cells from postcapillary venules of rat trachea overlap^11–13^. However, the characteristics and role of these overlaps during TEM have not previously been explored.

The opening and closing of the endothelial gap to allow the leukocyte to cross is very rapid and takes no more than a couple of minutes. Monitoring the dynamics of endothelial membranes during paracellular diapedesis, requires very fast image acquisition and high Z resolution. We used resonant confocal laser scanning and lattice light sheet microscopy to study leukocyte TEM in 3D and real-time and found that overlapping endothelial membranes form structures that we have termed “TEM tunnels” during diapedesis.

Recent work from our group showed that when endothelial membrane overlaps were artificially increased, using light-induced recruitment of the RacGEF Tiam1 to the membrane to locally activating the small GTPase Rac1, a rapid increase in the electrical resistance of endothelial monolayers is measured^35^. Additionally, when inducing a junction membrane protrusion (JMP), in only one of the two adjacent endothelial cells, we found these JMPs to function as TEM hotspots for leukocytes^28^.

A picture emerges that suggests that TEM events occur at areas of high endothelial membrane dynamics: the endothelial cell on top of the overlap shows increased protrusion activity, recognized by the crawling leukocyte, followed by a continuation of migration of the leukocyte on the bottom endothelial cell, hence, generating an endothelial-based membrane tunnel that supports leukocyte crossing. Interestingly, and in line with the idea that TEM events prefer high EC membrane dynamics, increased EC membrane flaps, supporting an exaggerated and faster level of neutrophil TEM (e.g. formation of hot-spots) is a feature of autophagy-deficient ECs^36^. As compromised autophagy is a hallmark of ageing and numerous inflammatory conditions, these findings offer a pathophysiological setting under which the presence of EC membrane flaps is pronounced.

The formation and stability of membrane overlaps do not depend on any of the junctional proteins investigated here, PECAM-1 and VE-cadherin. Whereas PECAM-1 is found to decorate the TEM tunnel, VE-cadherin is excluded from the tunnel and displaced to the sides. After the passage of the leukocyte, the two endothelial membranes connect again, and VE-cadherin re-distributes to the junction. It was already suggested that a maximized contact surface between endothelial cells and leukocytes would limit vascular leakage during TEM^4^. This offers another argument for the preference of transmigrating leukocytes for wide overlapping junctions. Indeed, we observed that leukocytes prefer transmigrating through wider PECAM-1-positive junctions *in vitro* and *in vivo*, underscored by our previous work and that of others^34,36^.

This study is the first to focus on endothelial membrane dynamics during leukocyte TEM at high spatiotemporal resolution to discover TEM tunnels during paracellular migration. Although LLSM is currently state-of-the-art to measure fast biological processes in real-time in Z, image processing, which includes deskewing, deconvolution, and volume rendering, is still required due to the angled light sheet illumination. Additionally, during TEM, neutrophils disturbed the light sheet and created shadow-like patterns on the endothelium that could not be corrected. Neutrophils are highly granularized, as they contain many inflammatory mediators in vesicles that can be secreted when pathogens are encountered. Unfortunately, these structures caused small deflections in the light sheet. We were not in the position to use other leukocyte types that are known to have less granules, e.g., T-lymphocytes or monocytes. But such experiments may be carried out in the future.

Altogether, this study defines the multicellular architecture of paracellular transendothelial migration, providing a three-dimensional context in which findings on this process can be placed.

## Acknowledgement

Lattice light sheet microscopy was performed at the Advanced Imaging Center at the Janelia Research Campus of Howard Hughes Medical Institute, USA. We wish to thank Dr. Paul Kubes and Dr. Björn Petri for their valuable discussion. This work was supported by LSBR grant # 1649 (A.C.I.v.S.), LSBR grant # 1820 (L.K.), and ZonMW NWO Vici grant #91819632 (J.D.v.B., W.J.v.d.M). SN is funded by the Wellcome Trust (098291/Z/12/Z and 221699/Z/20/Z to SN), LR is supported by funding from the British Heart Foundation (FS/IBSRF/22/25121)

## Materials and methods

### Mouse lines

Mice carrying the Rosa26-Confetti transgene were bred with Cdh5-CreERT2 mice to induce endothelial-specific, tamoxifen-inducible conditional Cre-recombinase expression in endothelial cells resulting in the exclusive labelling of endothelial cells with the heterogeneous labeling of the confetti construct. These confettifl/wt -Cdh5-CreERT2 mice were injected for five consecutive days with Tamoxifen (2mg/mouse) to induce sufficient recombination of the rosetta construct for imaging purposes. To preserve fluorophore function and avoid the collapse of blood vessels mice were perfused with PLP-fixation buffer before sacrifice by cervical dislocation. Mice were put into deep anesthesia using hexafluorane and the thorax was opened after which a cut to the left ventricle was made. 10 ML PLP fixation buffer was slowly injected into the left ventricle replacing the blood. After this procedure, the mouse was sacrificed by cervical dislocation, and organs were harvested.

The *Lyz2-EGFP-ki* mouse line was generously supplied by Dr. Markus Sperandio from Ludwig Maximilians University of Munich, Germany, and utilized with the consent of Dr. Thomas Graf from the Center for Genomic Regulation and ICREA, Spain. These *Lyz2-EGFP-ki* mice have EGFP cDNA integrated into the lysozyme M (Lyz2) locus, resulting in the production of EGFP+ myeloid cells, including GFPbright neutrophils, and GFPdim monocytes and macrophages^37^.

### Animal tissue preparation and staining

Liver, lung, and spleen were further fixated overnight using 4% PFA in PBS, washed with P-buffer, and stored in 30% sucrose overnight. The organs were then put into cryo molds filled with Tissue-TEK, frozen, and stored at -80. Afterwards, coupes were sliced and mounted for imaging using Prolong antifade Glass. Imaging was performed on a Leica SP8 and image analysis was performed using Imaris. The macaque retinas were obtained as described^16^. In brief, animal eyes were kindly provided by TNO Rijswijk and local laboratories. Institutional guidelines regarding animal experimentation were followed. The anterior segment including the ciliary body and iris were dissected from these eyes and processed for transmission electron microscopy analysis.

### Cell culture and transduction

Pooled human umbilical vein endothelial cells (HUVECs; Lonza, P1052, #C2519A) were cultured at 37◦C with 5% CO2 in enriched Endothelial Growth Medium (EGM2; #C-22211, Promocell) supplemented with 2% endothelial growth factor mix (#C-39216, Promocell), 100 U/mL penicillin, and 100 µg/mL streptomycin (#15140122, Gibco). Cells were cultured on fibronectin (FN; CLB) coated surfaces for up to passage 8. For fluorescence microscopy, cells were transduced (1:500) with lentiviral constructs to express CAAX tagged with fluorescent protein mScarlet, YFP, mNeonGreen, or mTurquoise2, and selected with 100µg/mL puromycin in EGM2. Cells were allowed to form a confluent mosaic monolayer after mixing transduced cells 1:1 in the combinations mScarlet/mNeongreen- and YFP/mTurqoise2-CAAX to enable overlap quantification as described later. To mimic inflammation, confluent cells received 10ng/mL human TNF-α (Peprotech, #300-01A) in EGM2 4 hours prior to imaging.

### Sample fixation

For antibody stainings, mosaic monolayers of two-color-CAAX-transduced HUVECs were cultured at FN-coated 12mm diameter glass imaging coverslips. Cells were fixed with 4% final concentration paraformaldehyde in phosphate buffered saline ++ (PBS, Fresenius Kabi, #M090001/02) with 1µg/mL CaCl_2_ and 0.5 µg/mL MgCl2 for 5 minutes at 37^◦^C and treated with antibodies and/or dyes (as specified). Coverslips were secured on imaging slides using Mowiol mounting medium (Sigma-Aldrich).

### Cytoskeleton manipulation during live imaging

Confluent mosaic monolayers of two-color-CAAX-transduced HUVECs were cultured at FN-coated 8-well µ-slides (#80826, Ibidi). F-actin modulating compounds were added in warm EGM2 to live cells during imaging at 37◦C with 5% CO2 to a final concentration of 1µM Cytochalasin B (Sigma-Aldrich, #C26762), 10µM Y-27632 (Calbiochem, #688000), 100µM CK-666 (Sigma-Aldrich, #SML006), or 10mM DMSO (Sigma-Aldrich). Afterwards, cells were fixed with PFA as described above and endothelial cell overlap was quantified using ImageJ/FIJI as described below.

### Lattice Light Sheet Microscopy imaging

The lattice light sheet microscope located at the Advanced Imaging Center (AIC) at the Janelia Research Campus of the Howard Hughes Medical Insititute (HHMI)^24^ was used and described previously^28^. In brief, HUVECs stably expressing mNeonGreen-CAAX or mScarlet-I-CAAX were cultured on FN-coated 5mm round glass coverslips (Warner Instruments, Catalog # CS-5R) for 2 days. ECs were stimulated with 10 ng/ml TNF-alpha 20h before imaging. Imaging was performed in HEPES buffer^21^ at 37 Celsius with 5% CO2 for maximally 30 min. Neutrophils were isolated as described before/below, stained for 20 min at 37 Celsius with Cell Tracker Deep Red (Invitrogen), washed and centrifuged for 3 min, 400G at room temperature, and added right above the coverslip between the excitation and detection objectives. 488 nm, 560 nm, and 642 nm diode lasers (MPB Communications) at 30% acousto-optic tunable filter (AOTF) transmittance and 50 mW initial box power and an excitation objective (Special Optics, 0.65 NA, 3.74 mm WD) were used for illumination. Fluorescence was detected via the detection objective (Nikon, CFI Apo LWD 25XW, 1.1 NA) and a sCMOS camera (Hamamatsu Orca Flash 4.0 v2). Exposure time was 20 ms with 50% AOTF transmittance and Z-step size was 0.211μm. The time interval was about 7.5 s for three-channel, 5 s for two-channel time-lapse, and 2.5 s for one-channel time-lapse. Point-spread functions were measured using 200 nm tetraspeck beads (Invitrogen cat# T7280) for each wavelength. Data was deskewed and deconvolved as described in Supplemental Methods and analyzed using Imaris software.

### Neutrophil isolation

Polymorphonuclear neutrophils (PMNs) were isolated from whole peripheral blood from healthy donors. Blood was diluted (1:1) in RT PBS with 1:10 TNC, transferred onto 1.076g/mL Percoll separation medium at RT, and separated by centrifuging at RT for 20 minutes at 800G with start and brake at setting 3. The supernatant was removed to leave only Percoll containing the PMNs and the erythrocyte layer below. Erythrocytes were lysed in ice cold buffer (water for injection with 155mM NH4Cl, 10mM KHCO3, 0.1mM EDTA (all Sigma-Aldrich)) on ice for 5-15 minutes until the suspension either cleared or became dark red. Neutrophils were pelleted at 4◦C for 5 minutes at 450G with start and brake at setting 9. Supernatant was removed and remaining erythrocytes were lysed again with ice-cold buffer for 5 minutes on ice. Neutrophils were pelleted and supernatant discarded, after which neutrophils were washed with 4◦C PBS and pelleted again. Neutrophils were then resuspended in HEPES+ buffer (20mM HEPES, 132mM NaCl, 6mM KCl, 1mM MgSO4, 1.2mM K2HPO4, pH7.4, 1mM CaCl2, 5mM D-glucose (all Sigma-Aldrich) and 0.4% human serum albumin (Sanquin Reagents)).

### Neutrophil transendothelial migration under flow

Confluent mosaic monolayers of two-color-CAAX-transduced HUVECs were cultured at FN-coated 6-channel µ-slides VI 0.4 (#80666, Ibidi)To mimic inflammation, cells received 10ng/mL human TNFα in EGM2 4 hours prior to flow experiments. During microscopy at 37◦C and 5% CO2, channels were connected to a pump system providing a laminar flow of 0.5dyne/cm2 in 37◦C HEPES+ medium. Every channel received 1 million freshly isolated neutrophils resuspended in HEPES+ medium that were activated by incubation at 37◦C for 15-30 minutes. Neutrophils were visualized using 1:6000 DiD dye that incubated with the neutrophils during activation. Transmigration was captured for 10 minutes after neutrophil injection to the flow system by acquiring repeated Z-stacks with step size never exceeding 1µm. Fluorescence was detected with the Zeiss LSM980 AiryScan2 machine (ZEISS) with ZEN software, using the 25x water-immersion NA.8 objective at 2.5x zoom and 8x multiplex imaging settings. Endothelial cell junction overlap was quantified using ImageJ/FIJI as described, classifying junctions based on presence or absence of a neutrophil transmigration event.

### Antibodies

Antibodies were used on fixed cells (see above) that were permeabilized, if necessary, 0.1% TritonX for 5-10 minutes. Cells were blocked using PBS with 2% bovine serum albumine (SERVA, #11920) and incubated with primary and secondary antibody, washing coverslips with PBS++ in between permeabilization, blocking and staining steps. Antibodies and dyes used: VE-cadherin mouse monoclonal IgG1 antibody directly labeled with Alexa Fluor 647 (BD Biosciences, #561567), PECAM-1 non-blocking mouse monoclonal IgG1 antibody directly labeled with Alexa Fluor 647 (BD Biosciences, #561654), JAM-A rabbit polyclonal IgG antibody (Zymed, #36-1700), Phalloidin labeled with Fluorescence dye 405-I (Fisher Scientific, #16109370), Hoechst 33342 (Molecular Probes, Life Technologies, #H-1399), Chicken anti-rabbit IgG secondary antibody Alexa Fluor 647 (Invitrogen, #A12443). Anti PECAM-1 for western blot, rabbit polyclonal IgG (Santa Cruz, #sc-8306), secondary swine anti-Rabbit-HRP antibody (DAKO, #P0399).

### In vitro Imaging

Live cells were imaged at 37◦C and 5% CO2. Overlap was visualized by acquiring Z-stack images using fluorescence microscopy, Z-step size never exceeding 1µm. On the Leica SP8 machine with LAS X software (Leica Microsystems B.V.) images were acquired at 1024*1024 resolution using the 40x NA1.4 oil-immersion objective and fluorescence was detected using PMT and HyD detectors with suitable gain and AOBS filter settings. On the Zeiss LSM980 AiryScan2 machine (ZEISS) with ZEN software the 40x NA1.4 oil-immersion objective was used for imaging at near-superresolution at 1.5x Nyquist-sampling. On both microscopes, excitation lasers were 405nm for Hoechst and Alexa405, 442nm for mTurqoise2, 488nm for mNeonGreen, 514nm for YFP, 561nm for mScarletI, 633nm for DiD, 633 nm and 671nm for respective Alexas.

### Confocal overlap analysis

To quantify overlap, maximum projections of Z-stacks were obtained using ImageJ/FIJI. Before thresholding, noise was reduced using the despeckle tool and by using the 2D median filter with radius 2.0 pixels. Fluorescent vesicles were removed using the particle analysis tool or by hand. EC overlap width, length and area were quantified using ImageJ/FIJI measure tool.

### Quantification of travel distance

PMN travel distances were quantified between first adhesion of the PMN on the monolayer and completed diapedesis. The travel start point was considered the PMN center of gravity upon adhesion. Then, the PMN center of mass was followed as it traveled the endothelium. The end point was considered the outer end of the overlap at the transmigration location.

### In vivo imaging of TEM tunnels

Ischemia reperfusion injury (IR) was performed according to^36^. Briefly, AF-conjugated non-blocking anti-PECAM-1 mAb (4 μg in 400 μL PBS) was administered by intrascrotal (i.s.) injection for 2 h. Cremasteric IR injury was then induced in the exteriorized cremaster muscle of anaesthetized mice by placing two non-crushing metal clamps (Interfocus, Schwartz Micro Serrefine) at the base of the exteriorized tissue (40 min). Subsequently, the clamps were removed to allow correct reperfusion of tissue and immunostained post-capillary venules (diameter: 20-40 μm) of the cremaster muscle were imaged using an upright Leica TCS SP5, with argon and helium lasers (488, 561 and 633 nm excitation wavelengths), using water dipping 20x/1.0 objective lens. TEM events were recorded by taking serial Z-stacks for 1-2 hour duration. A half-blood vessel was recorded, and the merged stack file/video represented an “en-face” view of a selected post-capillary venule. Imaris software was used to apply an isosurface of PECAM-1 to image the TEM tunnel.

### Statistical analysis

Data was plotted and analyzed using Prism software. Statistical tests used as indicated.

